## Supplementary material for "Centriole growth is not limited by a finite pool of components, but is limited by the Cdk1/Cyclin-dependent phosphorylation of Ana2/STIL": Reviews from Review Commons

Reviewer #1 (Evidence, reproducibility and clarity (Required)):

Centriole duplication is a conserved pathway that need to be tightly regulated. The key enzyme of centriole assembly is Plk4 which is recruited to the centrioles and undergoes dynamic re-localization from a ring-like pattern around a centriole to a dot-like morphology at the daughter centriole assembly site. This event is central for inducing centriole biogenesis. Plk4 then phosphorylates Ana2/STIL which allows recruitment of Sas-6 to form the cartwheel structure for centriole assembly.
In the present study, Steinacker, Wong et al. monitor how cytoplasmic concentrations of the key proteins in centriole assembly, Plk4, Asl/Cep152, Ana2/STIL, Sas-6 and Sas-4/CPAP change during the centriole assembly process in the Drosophila embryo by using fluorescence correlation spectroscopy (FCS) and Peak Counting Spectroscopy (PeCoS). They find that their concentrations remain constant with exception of Ana2/STIL of which cytoplasmic diffusion rate increased at the end of S-phase and is dependent on phosphorylation by Cdk1/CyclinB. Phosphorylated Ana2/STIL blocks centriole duplication thus preventing premature initiation of centriole duplication in mitosis.

Major comments
The manuscript is interesting and very well written. Most of the experiments are carefully performed. However, there are some important aspects for improvements that are listed below

Additional experiments:
- Figure 3: the transgenic flies that were generated here, ∆CC and ∆STAN, still contain wild-type Ana2. So, the authors therefore need remove or dampen their claim that the change in Ana2's cytoplasmic diffusion does not depend on its interaction with Sas-6 (page 11).
- Figure 5A: is the observed reduced recruitment of Sas-6 by Ana2(12A) due to a decrease in binding affinity? This should also be shown by analyzing protein-protein interactions between Ana2(12A) and Sas-6 biochemically.
- The authors use an Ana2(12A) mutant which comprises putative Cdk1 phosphorylation sites that have been identified in Mc Lamarrah et al. JCB 2018. However, only three of them were phosphorylated by Cdk1/cyclin B in vitro (Fig. S6). Are all these 12 putative Cdk1 phosphorylation sites important in vivo? Did the authors generate the Ana2(3A) or the S284A/T301A mutants to see whether it can rescue the ana2-/- mutant phenotype similar to the 12A mutant? These might be sufficient to observe the phenotype.
- Figure 6: is the interaction between Plk4 and Ana2(12A) impaired? Similarly, Plk4 activity and phosphorylation of Ana2(12A) by Plk4
- Figure 7: Phosphomimetics, in this case 12 amino acid changes, have the disadvantage of introducing more negative charge than the phosphorylated residue. The Ana2/(12D/E)-mNG is not efficiently recruited to centrioles. Is effect also observed for the Ana2/(3D/E) mutant?

Minor comments
Figure S1: only mNG-tagged centriolar proteins are shown. An empty mNGtag or an mNG-tagged non-centriolar protein should be shown to exclude that the tag by itself shows centriolar localization or somehow affects the localization
S4C: Sas6-mNG CPM error bars are missing for the 10min time point
S5A: What are the expression levels of the Ana2(12A) mutant? The expression levels shown in this Figure are not similar.

Reviewer #1 (Significance (Required)):

Centriole duplication normally begins at the G1/S phase transition. An important question in

the field is how premature centriole duplication in mitosis is prevented. The authors used fluorescence correlation spectroscopy (FCS) and Peak Counting Spectroscopy (PeCoS) to study the major conserved proteins in the centriole assembly pathwayq and found that only Ana2/STIL's cytoplasmic diffusion increases at the end of S-phase. It is known from the literature that Cdk1 prevent Plk4-STIL complex assembly in centriole biogenesis by directly competing with Plk4 for the CC domain of Ana2/STIL (Zitouni et al. Curr Biol 26, 1127-1137 (2016). However, Ana2/STIL can also bind to Plk4 via its conserved C-terminal region of STIL (Ohta et al., Cell Reports 11, 2018; McLamarrah et al., J Cell Biol 2018, 217, 1217-1231). The work by Steinacker, Wong et al. suggest that at least in fly embryos, growth of the daughter centriole is regulated though phosphorylation of Ana2 by Cdk1/CyclinB rather than binding.
The findings described in this manuscript are interesting for a broad range of scientists from both the centrosome and mitosis fields

Expertise of the reviewer: centriole biogenesis, structural and numerical centrosomal aberrations in disease

Reviewer #2 (Evidence, reproducibility and clarity (Required)):

Centriole growth is not limited by a finite pool of components, but is limited by the Cdk1/Cyclin-dependent phosphorylation of Ana2/STIL
Authors: Thomas L. Steinacker, Siu-Shing Wong, Zsofia A. Novak, Saroj Saurya, Lisa Gartenmann, Eline J.H. van Houtum, Judith R. Sayers, B. Christoffer Lagerholm, Jordan W. Raff
Centriole biogenesis is a tightly regulated process that occurs once per cell cycle. Defects in this process can lead to the acquisition of abnormal centriole numbers which has been linked to several human diseases. Centriole duplication starts with the assembly of a procentriole on the mother centriole in early S-phase followed by procentriole growth during G2 phase. A big question in the centrosome field is how new procentrioles assemble at the right time and acquire the correct final size.

In this manuscript, Wong et al. analyse whether the cytoplasmic concentration of several proteins changes during centriole assembly (Asl, Plk4, Ana2, Sas-6, and Sas-4). The authors show that the cytoplasmic concentration of these proteins remains constant during centriole duplication, indicating they are not limiting components for procentriole assembly. Nevertheless, the authors found that Ana2/STIL's cytoplasmic diffusion rate increases before the onset of mitosis, concurrent with an increase in Cdk1/Cyclin activity. Mutation of 10 putative phosphorylation sites in Ana2 prevented the diffusion rate change and enabled centrioles to grow for a longer period. This suggests that phosphorylation of Ana2/STIL by Cdk1/Cyclin could control the period of centriole growth.

Minor points:
In the introduction, the authors describe how PLK4 is required to recruit STIL and Sas-6 to promote the formation of the cartwheel during centriole duplication. However, there is also literature describing a role for STIL in regulating PLK4 abundance and localization pattern (i.e ring or dot) at the centriole.

The authors note that the levels of the Ana2(12A) mutant keep increasing until the onset of mitosis. The authors claim that this phenotype is consistent with the timing of increased Cdk1 activity. It would be interesting to show the increase in Cdk1 kinase activity over the same time-course and test whether dampening Cdk1 has the same effect on Ana2 recruitment.

While I appreciate detecting in vivo phosphorylation sites can be very challenging, It would be valuable to show the 10 Ana2 phosphorylation sites can be phosphorylated by Cdk1, at least in vitro.

Other points:

Figure 3: Amino acids numbers for CC domain are not the same in the figure and in the figure legend.

Figure 5Aii, the x-axis should be changed to minutes for easier comparison with other figures.

There are some typos in the figure legends.

Reviewer #2 (Significance (Required)):

This study attempts to address a central question in the centrosome field: how centriole growth is controlled. Although the paper does not provide a detailed mechanistic advance, the authors do provide some evidence against a limited pool of centriole components controlling centriole length, and they are careful not to overstate conclusions. The manuscript is well written and easy to follow. While it is not clear at present how phosphorylation of Ana2 alters its diffusion rate or limits centriole growth, I feel the study will be of interest to members of the centriole community and will stimulate new lines of investigation. Given that the cartwheel stops elongating in S phase in mammalian systems, it is not clear if the mechanism proposed would be conserved. That notwithstanding, I found this to be a rigorous study that advances our understanding of the regulation of the centriole duplication.

Reviewer #3 (Evidence, reproducibility and clarity (Required)):

The manuscript entitled "Centriole growth is not limited by a finite pool of components, but is limited by the Cdk1/Cyclin-dependent phosphorylation of Ana2/STIL" by Steinacker et al. nicely demonstrates that centriole growth in Drosophila embryos is not limited by a finite pool of core centriole components as in other systems. In contrast, they unveiled a specific elevated cytoplasmic diffusion rate of Ana2/STIL towards the end of the S-phase, correlating with the rise of Cdk1/Cyclin activity, that they hypothesize is important for the abrupt stop of centriole growth before mitosis (end of S phase). They found using an Ana2 mutant (12A) that cannot be phosphorylated by Cdk1/Cyclin that this elevated diffusion rate is abrogated, demonstrating that this kinase is involved in this process. The authors further conclude that daughter centrioles grow at a slower rate for an extended period (as followed by SAS-6 incorporation at centrioles in the context of the 12A mutant). Thus, the authors conclude that this novel mechanism ensures that daughter centrioles stop growing at the correct time and propose that it could be part of the explanation why centriole duplication does not occur during mitosis.
Overall, this is a solid study that is well written and easy to follow. The text and figures are well presented and the quality of the data is convincing. This manuscript would be of great interest not only to the centrosome field but also more generally to cell biologists.
I do not have major concerns regarding the experiments. However, I would like to propose some minor comments/clarification in order to further improve the manuscript.

Suggestions for additional improvements:

My main comments are related to the phosphorylated mutants of Ana2 (12A) and (12D/E).
1) To study the impact of Cdk1 on Ana2, the authors generated a mutant where 12 potential Cdk1 sites have been replaced by Alanine (12A). Although I acknowledge that all controls were properly done on this mutant and that the 12A protein is functional since it rescues the ana2-/- mutant phenotype, one can still wonder whether this could not affect somehow the overall protein conformation, or structure. Maybe this could simply be stated somewhere in the manuscript.

2) The authors mentioned that there is evidence that 10 Cdk1 sites in Ana2 are phosphorylated in vivo and they further demonstrate convincingly that 2 of the most conserved sites can be phosphorylated in vitro by Cdk1/CyclinB (Figure S6). Could the authors include the alignment showing the potential phosphorylation residues and highlight the 12 that were mutated and show the overall conservation of these sites? It would be easier to find the residues as from the scheme of Fig. 3A it is not easy to find which residues are mutated (although the information can be found in the method section p. 32).

3) My main confusion regarding the phosphorylation mutants 12A or 12D/E comes from the fact that both can rescue the ana2-/- mutant phenotype, which indicates that the mutant protein is functional and that somehow these sites are not fully important for centriole duplication or are not solely responsible for this type of regulation. Is this interpretation correct, this is somehow what I take from the end of the discussion p.21? if true maybe it should be a bit more emphasized.
4) Moreover, I would have expected since centriole growth does not stop abruptly (one could talk about "prolonged" centriole growth) in the 12A mutant that centrioles would be longer. However, this is not the case as shown in Figure S7. One possible explanation would be that even though the centriole growth is extended (looking at SAS6 as a proxy), the slope/rate of incorporation is lower. Could you please comment on this more? I think this is an important point of discussion/interpretation of the results.

4) The authors nicely show that the 12A mutant, despite similar expression levels as the taggedAna2WT, continued to accumulate at centrioles till NEBD (consistent with the hypothesis that Cdk1 cannot phosphorylate it and thus stops its recruitment). But how can the 12A levels decline at centrioles in mitosis where Cdk1 activity is the highest? This would mean that Cdk1 activity/level regulates Ana2 differentially over time or that other mechanisms might be at play. The authors mention in the discussion the attractive hypothesis of the "rheostat" (p. 20) but maybe a further discussion on an alternative mechanism could be also interesting.
Could the fact that the 12A level decreases in mitosis also explain the lack of centriole phenotype if we would imagine that levels at centrioles would stay high? Could the authors comment on this? They mention it briefly (p.14 and p.21) but if they could expand a bit would be great.

5) I was a bit confused about how the 12D/E mutant that is not recruited efficiently to centrioles could rescue the ana2-/- mutant centrioles? Could the authors comment on this, please?

6) p.16 still about the 12D/E that is not properly recruited to centrioles even in presence of one WT copy of Ana2 (untagged): The authors conclude that "phosphorylation at one or more of these S/T sites inhibits, but does not completely block, Ana2 recruitment to and/or maintenance at centrioles". Could the mutation also prevent Ana2 homo-oligomerization? In other words, could this result suggest that the 12D/E cannot interact with untagged Ana2WT and be recruited to centrioles? Is it a possibility?

Reviewer #3 (Significance (Required)):

In this manuscript, the authors address two major fundamental questions:
1- the mechanism that restricts strict cell cycle regulation of centriole duplication
2- How daughter centrioles grow to the correct size.
These questions are very important and this study provides some clues on the mechanisms that can be at play, among which Cdk1/cyclin seems to be involved.

In addition, this paper raises an interesting point in showing that the core centriole duplication components concentration is as low in human cells as in fast-dividing Drosophila embryos in the range of 5-20nM. This is very interesting as it was commonly thought that embryos would have a stockpile of core components to ensure fast and numerous centriole duplication cycles. Furthermore, they found that these concentrations remained constant using FCS or Pecos, demonstrating that core centriole components concentrations are not rate-limiting for centriole duplication (over time) in this system. Instead, they propose an alternative hypothesis whereby Cdk1/Cyclin phosphorylation of Ana2/STIL would be important to regulate centriole growth and ensured timely duplication (ie no duplication in mitosis, when Cdk1 activity is high).

In this context, this study would certainly have a broad interest and impact on cell biologists.

Reviewer's expertise:
Centrioles, microtubules, microscopy, cell biology.
