## Supplementary material for "Centriole growth is not limited by a finite pool of components, but is limited by the Cdk1/Cyclin-dependent phosphorylation of Ana2/STIL": Response to Reviews

**Manuscript number:** RC-2022-01297
**Corresponding author(s):** Jordan, Raff

### 1. General Statements [optional]

We would like to thank the reviewers for their helpful and constructive comments.

### 2. Point-by-point description of the revisions

**Reviewer #1**

This reviewer thought our findings would be of interest to a broad range of scientists from both the centrosome and mitosis fields, but noted some important aspects for improvements.

Additional Experiments (we number these points for ease of discussion).

*1. Figure 3. The reviewer points out that because our analysis of Ana2-∆CC and Ana2-∆STAN mutant proteins was conducted in the presence of endogenous WT protein, we should be more cautious in our interpretation.* We agree and apologise for overstating these findings. We have now rewritten the title and text of this section to be more cautious (p11, para.2)

*2. Figure 5A. The reviewer wonders whether the reduced recruitment of Sas-6 in the presence of Ana2(12A) is due to reduced binding, and they request we test this biochemically.* This is our favoured interpretation, but we have been unable to test this biochemically for two reasons. First, although we have successfully purified several recombinant Sas-6 and/or Ana2 fragments (Cottee et al., *eLife*, 2015), the full-length proteins are poorly behaved (tending to precipitate, likely due to their inherent ability to self-oligomerise). Thus, we have been unable to reconstitute their interaction *in vitro*. Second, as we show here, the proteins are normally expressed in embryos at surprisingly low concentrations (~5-20nM), and we can detect no interaction between them in coimmunoprecipitation experiments from embryo extracts (not shown). Indeed, this concentration is so low that Sas-6 does not even appear to form a homo-dimer in the embryo, even though Sas-6 clearly functions as a homo-dimer in centriole assembly (new Figure S4A). We now explain these points, and state that our favoured hypothesis that Ana2(12A) has reduced affinity for Sas-6 (or other core duplication proteins) remains to be tested (p22, para.2).

*3. The Reviewer wonders if all 12 of the potential Cdk1 phosphorylation sites that we mutate in Ana2(12A) are important in vivo, and whether we have tested whether mutating fewer sites (e.g. the two sites [S284/T301] that we show are phosphorylated by Cdk1/Cyclin B in vitro) might be sufficient to recapitulate the Ana2(12A) phenotype.* We have now tested this by mutating just the S284/T301 sites to Alanine [Ana2(2A)], but the results were not very informative (Reviewer Figure 1 [RF1]). Whereas Ana2(12A) is recruited to centrioles for a longer period and to higher levels than WT Ana2 (Figure 4A), Ana2(2A) is recruited to centrioles for a normal period but to lower levels (RF1A,B). The interpretation of this result is complicated because western blots show that

Ana2(2A) is also present at lower-levels than normal (RF1B). Thus, it is clear that Ana2(2A) does not recapitulate well the behaviour of Ana2(12A). We have decided not to present this data as it is difficult to interpret and it does not change any of our conclusions.

*4. Figure 6. The reviewer asks whether the 12A mutations impair the interaction with Plk4, influence Plk4's kinase activity or the ability of Plk4 to phosphorylate Ana2.* These are excellent questions but, for the same reasons described in point 2 above, we cannot address them biochemically as we cannot purify well-behaved recombinant full-length Ana2 or active Plk4 *in vitro*, and both proteins are present at such low levels in the embryo that we cannot detect any interaction between them in embryo extracts. We are working hard to reconstitute *in vitro* systems to probe these important points, but it may be sometime before we are able to do so.

*5. Figure 7. The reviewer suggests that the 12D/E phosphomimetic substitutions introduce more negative charge than the putative phosphorylation of Ser/Thr residues and they ask if the Ana2(2D/E) [stated as Ana2(3D/E)] is, like the Ana2(12D/E) mutant, not efficiently recruited to centrioles.* This is a fair comment, but we have not analysed an Ana2(2D/E) mutant because, as described in point 3 above, the Ana2(2A) mutant did not recapitulate well the Ana2(12A) phenotype.

Minor comments

*1. Figure S1. The reviewer requests that we show that the mNG tag on its own is not recruited to centrioles.* We do not show this (as it would create a lot of white space in this Figure), but now state that mNG and dNG do not detectably localise to centrioles (p7, para.1).
*2. Figure S4C.* We have included the missing error bars (now Figure S4B).
*3. Figure S5A. The reviewer asks about the expression levels of the Ana2(12A) mutant, which are not shown in this Figure. They also state that the expression levels of the transgenes shown in Figure 5A are not similar.* The expression level of Ana2(12A) is shown in Figure S9, as this data was analysed independently of the other mutant proteins shown in Figure S5. We agree that it was overly simplifying the situation to state that the expression levels of WT Ana2-mNG, eAna2(∆CC)-mNG and eAna2(∆STAN)-mNG were "similar" (Figure S5), and we now specifically mention the differences between them (p11, para.3).

**Reviewer #2**
This reviewer found this a rigorous study that advances our understanding of the regulation of centriole duplication, but raised some minor points.

Minor Points

*The reviewer requests that we mention the literature describing how Ana2/STIL can influence the abundance and centriolar localisation of Plk4.* We apologise for this omission, and have amended our description of this literature in the Introduction to include this point (p3, para.2).

*The reviewer notes that we interpret the ability of the Ana2(12A) mutant to keep incorporating into the centrioles for a longer period as being consistent with our idea that rising levels of Cdk activity during S-phase normally reduce the ability of WT Ana2 to bind to the centriole. They ask us to show how Cdk activity increases over this time-course, and to test whether dampening Cdk has the same effect on Ana2 recruitment (i.e. allows Ana2 to be recruited for a longer period).* The time-course of Cdk activation in these embryos has been reported previously (Deneke et al., *Dev. Cell*, 2016; we present the relevant data from this paper in RF#2A [*black line*]). This reveals how Cdk activity rises throughout S-phase, which is crucial for our model. To assess the effect of dampening Cdk activity in these embryos we have now analysed the effect of halving the genetic dose of Cyclin B (RF#2B). This perturbation extends S-phase length, but has a complicated effect on the recruitment dynamics of Ana2 (RF#2B). As we would predict, Ana2 is recruited to centrioles for a longer period in these embryos, but it is also recruited more slowly (so it accumulates to lower levels). This is consistent with our hypothesis that Cdk1 activity might first stimulate and then ultimately inhibit the centriolar recruitment of Ana2. The interpretation of this experiment is not straightforward, however, as dampening Cdk1 activity alters Ana2 recruitment dynamics (and many other processes in the embryo) in complicated ways, so we have decided not to include it in the manuscript.

*The reviewer suggests that it would be valuable to show that all 12 of the potential Cdk1 phosphorylation sites in Ana2 can be phosphorylated by Cdk1 in vitro.* We think this would not be particularly informative as our hypothesis does not rely on all 12 sites being phosphorylated to generate the Ana2(12A) phenotype. We simply mutate all 12 sites because we don't know which, if any, are relevant. Thus, showing that some/all of the 12 sites can/cannot be phosphorylated *in vitro* does not test any hypothesis and would not change any of our conclusions. We now explain our thinking on this in more detail (p12, para.2)

Other points
*Figure 3.* We have corrected the amino-acid numbering mistakes.
*Figure 5Aii.* We have changed the x-axis (time) labelling in this and all other Figures.
*Figure Legends*. We have tried to eliminate the typos from the Figure legends, and apologise that these errors made it through to the final submitted version of our manuscript.

**Reviewer #3**
This reviewer thought our manuscript would be of great interest to not only the centrosome field but also to cell biologists more generally. Although they had no major concerns, they made a number of suggestions for improvements.

*1.* As the reviewer suggests, we now explicitly state that although the Ana2(12A) mutant appears to be largely functional, the overall conformation of the protein may be altered, changing its function in ways we do not appreciate (p21, para.2).

*2. The reviewer suggests we include a multiple sequence alignment of Ana2/STIL proteins to provide more context about the distribution and conservation of the 12 S/T-P sites mutated in Ana2(12A).* This is an excellent idea, and we now include this in a new Figure S6, where we also provide more information about which of these sites have been shown to be phosphorylated in embryo or S2-cell extracts

*3. The reviewer is confused as to why the 12A and 12D/E mutants rescue the ana2$^{-/-}$ mutant flies so well, which suggests that the mechanism we propose here cannot be essential for centriole duplication.* We understand this confusion and we now make this point more clearly and explain why we think this occurs in more detail (e.g. p22, para.1). We propose that Cdk normally phosphorylates Ana2 to inhibit its ability to promote centriole duplication, but this phosphorylation does not entirely block this function. So, if all other elements of the system are functional, Ana2(12A) is recruited to centrioles for longer than normal, but this does not dramatically perturb centriole duplication because the many other factors that regulate centriole duplication (such as the pulse of Plk4 recruitment to centrioles [Aydogan et al., *Cell*, 2020]) still occur normally and are sufficient to ensure that centrioles still duplicate normally. When Ana2 phosphorylation is mimicked [Ana2(12D/E)], the ability of Ana2 to promote centriole duplication is perturbed (but not abolished). This perturbation is lethal in the early embryo—where the centrioles must duplicate in just a few minutes to keep pace with the rapid nuclear divisions. In somatic cells S-phase is much longer, so these cells can still duplicate their centrioles (as we observe) even though Ana2(12D/E) does not function efficiently. As we now explain, this phenotype (being lethal in the early embryo, but not in somatic cells) is a common feature of mutations that influence the *efficiency* of centriole and centrosome assembly (p17, para.2).

*4A. The reviewer asks us to comment in more detail on why centrioles do not seem to be elongated in the Ana2(12A) mutant wing disc cells (now Figure S8C), even though we show that Ana2(12A) (Figure 4A), and also Sas-6 (Figure 5), are recruited to centrioles for an abnormally long period.* This is an excellent question and, although we do not know the answer, we now discuss this interesting point in more detail (p16, para.1). We think this is likely due to the "homeostatic" nature of centriole growth: in our hands, almost any perturbation that makes centrioles grow for a longer/shorter period, also makes them grow more slowly/quickly, so that they tend to grow to a similar size (Aydogan et al., *JCB*, 2018; *Cell*, 2020). This is fascinating, but poorly understood. When we perturb the system by expressing Ana2(12A), both Ana2(12A) and Sas-6 incorporate into centrioles for a longer period, as we predict (Figure 4A and 5A). Unexpectedly, however, Sas-6 is also recruited to centrioles much more slowly. Thus, as so often happens, when we perturb the system so the centrioles grow for a longer time, the centrioles "adapt" by growing more slowly. We do not currently understand why this occurs (although we speculate that Ana2 may also be regulated by Cdk/Cyclins to help recruit Sas-6 to centrioles in early S-phase). In the embryo, where S-phase is very short, this homeostatic compensation is not perfect, and the centrioles appear to actually be shorter than normal. In somatic wing-disc cells, where S-phase is much longer, we suspect that there is more scope for homeostatic compensation and so the centrioles grow to the correct size.

*4B. In this point (also labelled [4] by the reviewer, so we have retained this numbering but labelled the points A and B) the reviewer asks why levels of Ana2(12A) eventually decline at centrioles once the embryos actually enter mitosis. The reviewer notes our rheostat theory, but suggests a discussion of other mechanisms might be interesting.* This is a good point, and we agree that the observation that Ana2(12A) levels ultimately still decline at centrioles during mitosis is likely to be important in explaining why centriole duplication is not more dramatically perturbed by Ana2(12A). We now expand our discussion of this point, highlighting that other mechanisms must help to ensure that Ana2 is not recruited to centrioles during M-phase, and discussing the possibility that the receptors that recruit Ana2 to centrioles are themselves inactivated during mitosis by high levels of Cdk activity (p15, para.1). In such a model, the rapid drop in WT Ana2 centriolar levels is due to a combination of switching off Ana2's ability to bind to centrioles (as we propose here) *and* switching off the ability of the centrioles to recruit Ana2. For Ana2(12A), only the latter mechanism would operate, so Ana2(12A) levels would start to drop later in the cycle (as the inflexion point at which Ana2 recruitment and loss balances out would be moved to later in the cycle), and these levels would drop more slowly—as we observe.

*5. The reviewer is confused to how the Ana2(12D/E) mutant can rescue the mutant phenotype when it is recruited to centrioles so poorly.* Ana2(12D/E) is indeed recruited very poorly to centrioles in the experiment shown in Figure 7. However, this experiment had to be conducted in the presence of WT untagged Ana2—as the embryos do not develop in the presence of only Ana2(12D/E). We would predict that WT Ana2 would bind more efficiently to centrioles than Ana2(12D/E) (which appears to behave as if it has been phosphorylated by Cdk/Cyclins, and so cannot be recruited to centrioles efficiently). Thus, in the experiment we show in Figure 7, the Ana2(12D/E) protein is probably being "outcompeted" for binding to the centriole by the WT protein. In somatic cells expressing *only* Ana2(12D/E) presumably sufficient mutant protein can be recruited to centrioles to support normal centriole duplication (as it no longer has to compete with the WT protein). We now explain our thinking on this point (p18, para.1).

*6. The reviewer wonders whether Ana2(12D/E) may be unable to homo-oligomerize, and this may explain why the protein is not recruited to centrioles efficiently even in the presence of WT protein.* This is indeed a possibility, but we think it unlikely as it is widely believed that Ana2/STIL proteins must multimerize to be functional (Arquint et al., *eLife*, 2015; Cottee et al., *eLife*, 2015; Rogala et al., *eLife*, 2015; David et al., *Sci. Rep.*, 2016). As Ana2(12D/E) strongly restores centriole duplication in *ana2*$^{-/-}$ mutant somatic cells, it seems unlikely that it cannot multimerize. Nevertheless, we now specifically highlight that the 12D/E (and 12A) mutations might alter the ability of Ana2 to multimerise (p21, para.2).

We thank the reviewers again for their thoughtful and constructive comments. We hope they will agree that the revised manuscript is now improved and would be appropriate for publication in *The Journal of Cell Biology*.

With best wishes,
